# Bandage: interactive visualisation of *de novo* genome assemblies

**DOI:** 10.1101/018333

**Authors:** Ryan R. Wick, Mark B. Schultz, Justin Zobel, Kathryn E. Holt

## Abstract

**Summary:** While *de novo* assembly graphs contain assembled contigs (nodes), the connections between those contigs (edges) are difficult for users to access. Bandage (a **B**ioinformatics **A**pplication for **N**avigating ***D**e novo* **A**ssembly **G**raphs **E**asily) is a tool for visualising assembly graphs with connections. Users can zoom in to specific areas of the graph and interact with it by moving nodes, adding labels, changing colours and extracting sequences. BLAST searches can be performed within the Bandage GUI and the hits are displayed as highlights in the graph. By displaying connections between contigs, Bandage presents new possibilities for analysing *de novo* assemblies that are not possible through investigation of contigs alone.

**Availability and implementation:** Source code and binaries are freely available at https://github.com/rrwick/Bandage. Bandage is implemented in C++ and supported on Linux, OS X and Windows.

**Supplementary information:** A full feature list and screenshots are available at *Bioinformatics* online and http://rrwick.github.io/Bandage.

## 1 INTRODUCTION

Current *de novo* genome assemblers use graphs, most typically a de Bruijn graph. An ideal graph would contain one distinct path for each underlying sequence, but complexities such as repeated sequences usually prevent this. Instead, assembly graphs contain branching structures, where one node may lead into multiple others. The longest sequences in the graph that can be determined unambiguously are saved as contigs, which are often the final result of *de novo* assembly (Schatz *et al.*, 2010).

The process of extracting contigs from an assembly graph involves discarding information; the connections between nodes and sequences too short to be useful on their own are not saved. It can however be advantageous to examine an assembly graph directly, where such connection information is clear.

Bandage facilitates interaction with de Bruijn graphs made by *de novo* assemblers such as Velvet (Zerbino *et al.*, 2008), SPAdes (Bankevich *et al.*, 2012) and Trinity (Grabherr *et al.*, 2011). It displays the graph in a graphical user interface (GUI) using a simple, comprehensible representation. The program is interactive, allowing users to zoom, pan and manually move nodes to focus on areas of interest.

## 2 IMPLEMENTATION AND PERFORMANCE

Bandage is a GUI application written in C++ with Qt, giving it speed, memory efficiency and cross-platform portability. It runs on Linux, OS X and Windows. The Open Graph Drawing Framework library (http://www.ogdf.net/) is used to perform the graph layout using the fast multipole multilevel layout algorithm, which scales well for very large graphs (Hachul *et al.*, 2007).

On a 3 GHz laptop, Bandage can load and display the *de novo* assembly graph for a bacterial genome (5 Mb) in a few seconds, using less than 100 MB of RAM. A large and complex graph, such a metagenome assembly, may take minutes and multiple gigabytes of RAM to display in its entirety, but the visualisation can be limited to smaller regions of the graph to improve performance and reduce memory requirements.

## 3 FEATURES AND CASE STUDIES USING ILLUMINA 100 BP PAIRED END READS

### 3.1 Assembly quality and completion

Assemblies of whole genomes can be difficult to complete if repeated sequences occur in chromosomes or plasmids. Repeated sequences cause distinctive structures in the assembly graph, limiting contig length. Bandage’s visualisation of the assembly graph makes it easy to identify such problematic parts of assemblies (Figure 1a). In some cases, it is possible to manually resolve these ambiguities by using additional information not available to the assembler. Bandage facilitates this by allowing users to copy sequences directly from the graph visualisation. In other cases, ambiguities cannot be resolved and the assembly cannot be completed. For these situations, Bandage provides a clear illustration of the assembly’s incompleteness and comparison of one assembly’s quality to another.

**Fig. 1.**
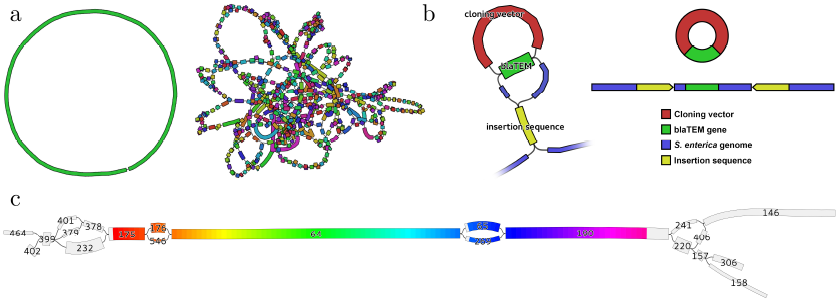
Examples of Bandage visualisation **(a)** Left, ideal bacterial assembly (single contig); right, poor assembly with many short contigs. **(b)** Left, zoomed-in view of Salmonella assembly; repeated sequences (blaTEM and insertion sequence) appear as single nodes with multiple inputs and outputs. Right, underlying gene structure deduced from Bandage visualisation. **(c)** 16S rRNA region of a bacterial genome assembly graph, highlighted by Bandage’s integrated BLAST search; node labels indicate node names, widths indicate coverage. Even though the 16S gene failed to assemble into a single node, the user can manually reconstruct a complete dominant gene sequence from this succession of nodes: 175, 176, 64, 65 and 190.

### 3.2 Resolving a complex antibiotic resistance region

Whole genome sequencing is frequently used to investigate evolution and transmission of bacterial pathogens and the mobile genetic elements responsible for antibiotic resistance. In this case study, multiple strains of *Salmonella enterica* were sequenced to determine the composition of their *Salmonella* genomic island (SGI), which can confer antibiotic resistance. The assembly of the SGI was complicated by sequences that were shared between the SGI, plasmids and a cloning vector used in the preparation of libraries for sequencing, making the contigs of limited use.

With Bandage, the user could search for nodes of interest (resistance genes or known SGI sequences) and zoom in to deeply investigate those areas of the graph. By showing the graph edges, Bandage makes clear which contigs are connected, their directionality and therefore which sequences are likely to be contiguous (Figure 1b). For many *Salmonella* strains, this allowed for determination that a gene of interest was or was not contained in the SGI, even when other methods were inconclusive.

### 3.3 16S sequence from a bacterial assembly

16S rRNA sequences are commonly used for classification of bacteria, but this is complicated by the fact that a single bacterial genome may contain multiple distinct copies of the 16S gene (Větrovský *et al.*, 2013). Some regions of the gene are highly-conserved while others regions are variable, potentially disrupting its *de novo* assembly.

In this case study, the user sequenced a bacterial isolate with whole genome shotgun sequencing and assembled the reads with Velvet v1.2.10 (Zerbino *et al.*, 2008). Bandage’s BLAST integration highlighted the 16S gene in the assembly graph (Camacho *et al.*, 2009). It did not assemble completely, and fragments of the gene were present in seven different nodes (Figure 1c). Consequently, the longest available 16S sequence in the contigs file was 909 bp and only partially covered the gene.

By illustrating graph connections and node coverage, Bandage made it clear which path represents the gene’s most abundant copy. The user was then able to extract node sequences from Bandage to manually reconstruct a 1535 bp sequence containing an entire 16S gene.

## 4 CONCLUSION

By visualising both nodes and edges, Bandage gives users easy, fast access to the connection information contained in assembly graphs. This is particularly useful when the assembly contains many short contigs – as is often the case when assembling short reads – and empowers users to examine and assess their assembly graphs in greater detail than viewing contigs alone.

## 5 ACKNOWLEDGEMENTS

I am grateful to both Simon Gladman and Jane Hawkey for their testing and feedback during Bandage’s development.

